# *Arabidopsis thaliana* phytocystatin 6 forms functional oligomer and amyloid fibril states

**DOI:** 10.1101/2023.09.25.559285

**Authors:** Naiá P. Santos, Hans Brandstetter, Elfriede Dall

## Abstract

Cystatins not only encode a high functional variability because of their ability to inhibit different classes of proteases but also because of their propensity to form oligomers and amyloid fibrils. Phytocystatins, essential regulators of protease activity in plants, specifically inhibit papain-like cysteine proteases (PLCPs) and legumains through two distinct cystatin domains. Mammalian cystatins can form amyloid fibrils, however, the potential for amyloid fibril formation of phytocystatins remains unknown. In this study, we demonstrate that *Arabidopsis thaliana* phytocystatin 6 (AtCYT6) exists as a mixture of monomeric, dimeric, and oligomeric forms in solution. Non-covalent oligomerization was facilitated by the N-terminal cystatin domain, while covalent dimerization occurred through disulfide bond formation in the interdomain linker. The non-covalent dimeric form of AtCYT6 retained activity against its target proteases, papain, and legumain, albeit with reduced inhibitory potency. Additionally, we observed the formation of amyloid fibrils by AtCYT6 under acidic pH conditions and upon heating. The amyloidogenic potential could be attributed to AtCYT6’s N-terminal domain (AtCYT6-NTD). Importantly, AtCYT6 amyloid fibrils harbored inhibitory activities against both papain and legumain. These findings shed light on the oligomerization and amyloidogenic behavior of AtCYT6, expanding our understanding of phytocystatin biology and its potential functional implications in plant protease regulation.

## Introduction

Amyloid fibrils are robust, insoluble protein polymers assembled in cross beta structures. The list of proteins reported to give rise to amyloid fibrils *in vivo* and *in vitro* reveals a heterogeneous landscape of protein families, native folds and functions across the kingdoms. Such diversity is observed whether they are involved in pathologies or are fibrils with known physiological functions [1–6]. Only recently, however, functional amyloids were described *in vivo* in Plantae. The garden pea (*Pisum sativum)* seed storage protein vicilin, a 7S globulin, was shown to form amyloids *in vivo* that arise during seed maturation and are consumed during germination [7]. Phytocystatins (phycys) belong to the cystatin superfamily, a fibril-forming family [8–13] of mostly small protease inhibitors that in plants are implicated in defense mechanisms [14–16]. Despite the significant amount of research on the fibril formation of human cystatins, there is currently no evidence that phytocystatins exhibit a similar tendency to form fibrils. Cystatins are potent, reversible competitive inhibitors of papain-like cysteine proteases (PLCPs) that fold into a characteristic antiparallel β-sheet wrapping a central α-helix. Phycys are functionally subdivided into 3 subfamilies. Type I phycys harbor one cystatin domain that inhibits PLCPs in an “elephant-trunk” binding mode, as opposed to a substrate-like mode. Specifically, three regions are responsible for PLCP binding: (i) an N-terminal segment preceding a conserved glycine, (ii) the L1 loop bearing the Q-X-V-X-G motif, and (iii) a P-W motif on the L3 loop. Type II phycys harbor two tandem cystatin domains that can inhibit legumain in addition to PLCPs, and type III phycys are large multidomain cystatins resulting from gene duplication [17]. The NMR structure of a type II phycys from *Sesamum indicum* (PDB 2MZV) illustrates the typical two-domain architecture that, differently from type II cystatins from humans, targets PLCPs via the N-terminal cystatin domain and legumains via the C-terminal cystatin domain [18, 19]. The legumain family of cysteine proteases are also known as vacuolar processing enzymes (VPEs) that show a strict specificity for asparagine or aspartic acid residues in P1 position. *A. thaliana* expresses four isoforms of legumain (AtLEG), namely AtLEGα, -β, -γ and -δ. AtLEGβ is found preferentially in seeds and is responsible for protein storage mobilization and pollen maturation [20, 21].

Phycys exploit the strict substrate preference of legumain and bind to it via a conserved asparagine residue on a reactive center loop that blocks its active site. This substrate-like interaction is further stabilized by charged interactions to an additional legumain exosite loop (LEL) on the phycys [22]. Importantly, cystatins are able to stabilize the pH-sensitive legumain in unfavorable pHs, as shown for human cystatin M/E (hCE) and phytocystatin 6 from *Arabidopsis thaliana* (AtCYT6) [12, 22]. Cystatins encode a high functional variability not only because of their ability to inhibit different classes of proteases, but also because of their propensity to form dimers, oligomers and amyloid fibrils. In human cystatins, dimers and fibrils are typically formed via a domain swapping (DS) mechanism upon conformational destabilization caused by pH shifts, mutations or truncations [12, 23]. In domain swapping, the N-terminal segment comprising β1-α1-β2 and the L1-loop of one monomer flaps out and repositions at the equivalent position of a second molecule and *vice versa* [24]. As a result, the L1 loop is disrupted, and consequently domain-swapped cystatin dimers are inactive as PLCP inhibitors. Importantly, their inhibitory activity against legumains, if present, remains unaffected by domain swapping [12, 24–26]. Furthermore, human cystatins M/E and C (hCC), stefin B and chicken cystatin were shown to form amyloid fibrils *in vitro* and *in vivo* via domain swapping [9, 10, 12]. Additionally, multiple members of the cystatin superfamily were shown to form DS-dimers, including phycys canecystatin 1 from sugarcane (*Saccharum officinarum*), cystatin 1 from hop (*Humulus lupulus*), and cystatin 1 from *Vigna unguiculata* [12, 27–30]. *Arabidopsis thaliana* is predicted to express seven phycys isoforms [31]. AtCYT6 is the only *A. thaliana* phycys with proven inhibitory activity against papain and AtLEGβ and -γ [22]. However, its propensity to form dimers, higher oligomers or amyloid fibrils was not studied so far. Therefore, we set out to characterize its conformational variability. Here we show that AtCYT6 forms amyloid fibrils *in vitro*, and that its individual cystatin domains feature strikingly different amyloidogenic potentials that could translate into a regulatory mechanism for PLCP and legumain inhibition.

## Results

### AtCYT6-NTD triggers oligomer formation within AtCYT6

To investigate if AtCYT6 forms amyloid fibrils, we designed, expressed, and purified to homogeneity four different AtCYT6 constructs: (i) full-length AtCYT6 (AtCYT6) harboring the N- and C-terminal cystatin domains, (ii) the isolated N-terminal domain (AtCYT6-NTD), (iii) the isolated C-terminal domain (AtCYT6-CTD) and (iv) the C-terminal domain containing the inter-domain linker at its N-terminus (AtCYT6-CTD_long_). Size exclusion experiments provided the first evidence that AtCYT6 formed oligomers in solution (Fig. 1A). At pH 7.5 and under non-reducing conditions, AtCYT6 eluted as a mix of at least four distinct oligomeric species, corresponding to the expected sizes of monomeric, dimeric and higher oligomeric forms. Cross-linking experiments using glutaraldehyde further confirmed the observed dimer (Fig. 1E). Since AtCYT6 harbors a single cysteine (Cys138) in the inter-domain linker connecting the N- and the C-terminal domain, we suspected that oligomerization could be mediated by disulfide formation. To test this hypothesis, we repeated the experiment under reducing conditions. As seen in Fig. 1A and S1A, AtCYT6 assembled into oligomers, in both conditions. However, overall, the peak profile shifted to the lower molecular weight monomer and dimer species under reducing conditions.

**Figure 1.**
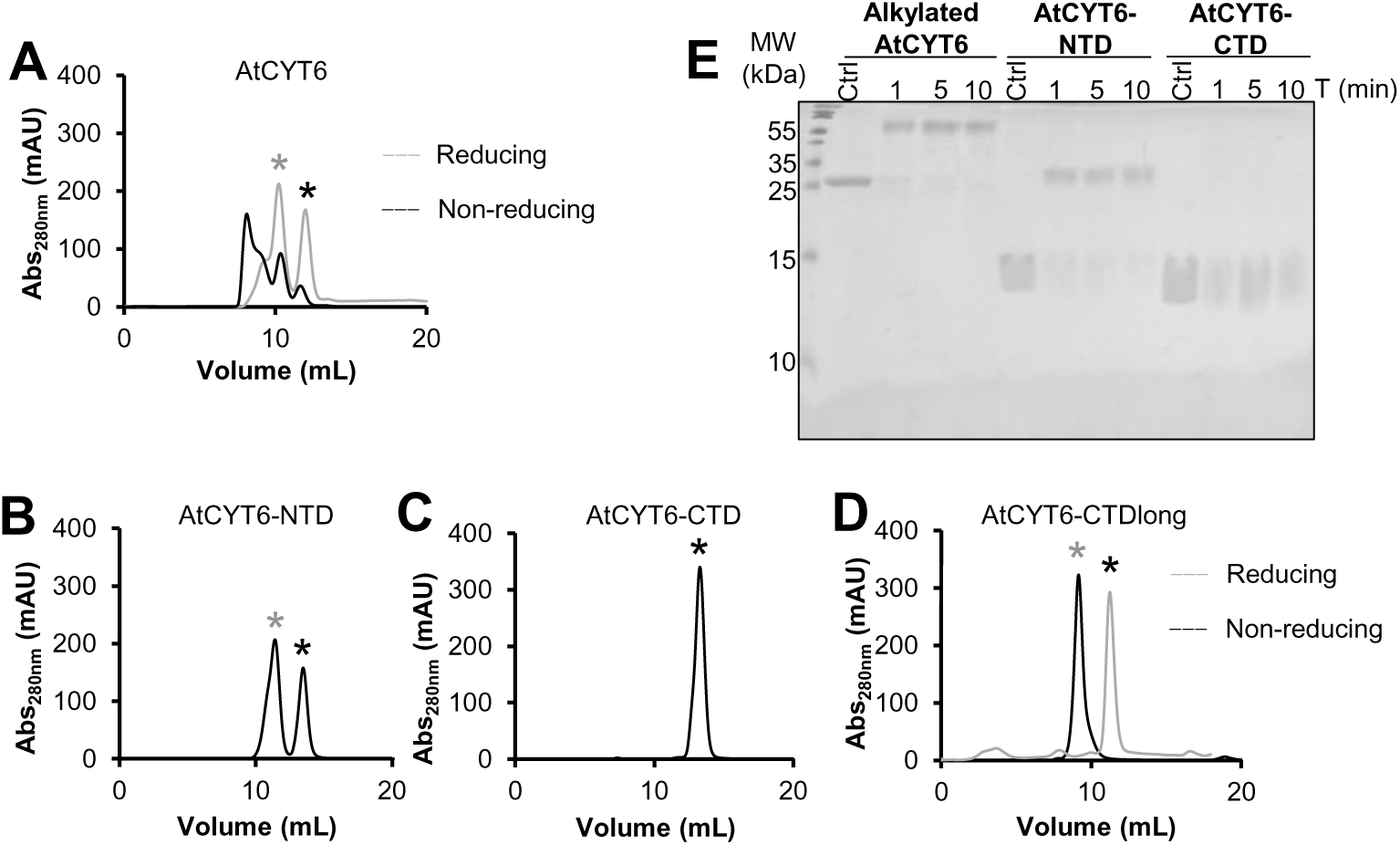
AtCYT6-NTD and Cys138 on the inter-domain linker mediate dimer and oligomer formation in AtCYT6. **(A)** AtCYT6 was subjected to size exclusion chromatography experiments at pH 7.5 in the presence (grey curve) or absence (black curve) of the reducing agent DTT (dithiothreitol). The expected elution volume of monomeric and dimeric AtCYT6 is indicated by a black and a grey star respectively. In the presence of DTT, high molecular weight oligomers were reduced to monomers and dimers, suggesting disulfide-mediated oligomerization via Cys138 on the inter-domain linker in addition to non-covalent dimer formation. **(B)** SEC profile of AtCYT6-NTD under non-reducing conditions (no cysteine is present in AtCYT6-NTD) revealed two peaks corresponding in size to monomeric and dimeric forms. **(C)** Same as (B) but using AtCYT6-CTD, which does also not harbor a cysteine in its sequence and eluted as a single peak corresponding in size to monomeric AtCYT6-CTD. **(D)** Same as (C) but using the AtCYT6-CTD_long_ construct which carries the inter-domain linker on its N-terminal end. Under non-reducing conditions AtCYT6-CDT_long_ migrated at the expected size of a dimer. The dimer peak resolved into a monomer peak in the presence of DTT, implying disulfide mediated dimerization via Cys138 on the inter-domain linker. **(E)** Cross-linking experiments with glutaraldehyde for 1, 5 and 10 minutes revealed that while AtCYT6 and AtCYT6-NTD were present as monomers and dimers in solution, AtCYT6-CTD was only present as a monomer. AtCYT6 was alkylated before setting up the experiment to prevent disulfide-mediated dimerization via Cys138.

When we analyzed the isolated N-terminal domain AtCYT6-NTD, we found that it eluted as a double peak corresponding to monomeric and dimeric species. Cross-linking experiments again confirmed the observed dimer (Fig. 1B,E). Importantly, AtCYT6-NTD did not harbor a cysteine residue in its sequence. This finding was in stark contrast to AtCYT6-CTD, which was also lacking Cys138 and eluted exclusively as a monomer in SEC experiments under non-reducing conditions (Fig. 1C). In line with this observation, AtCYT6-CTD did also not show dimer formation in cross-linking experiments (Fig. 1E). Interestingly, AtCYT6-CTD_long_, which contained an unpaired Cys138 on the N-terminal linker extension, eluted as a dimer in the absence of DTT (Fig. 1D). Upon addition of DTT, the dimer peak shifted to the expected molecular weight of monomeric AtCYT6-CTD_long_. Altogether, Cys138 was able to mediate covalent dimer formation of AtCYT6 *in vitro*, and AtCYT6-NTD is likely responsible for noncovalent dimer formation by AtCYT6.

### Modeling suggests that AtCYT6 dimers may form via domain swapping

Domain swapping is a hallmark of the oligomerization of human cystatin isoforms finally resulting in the formation of dimers and amyloid fibrils [8]. To gain insight into the structural flexibility of AtCYT6, we used AlphaFold Multimer to model different oligomeric states. AlphaFold provides a per-residue assessment of the confidence of the prediction with the predicted *local distance difference test* (pLDDT). A pLDDT value higher than 90 represents a *high confidence* prediction, and a pLDDT value higher than 70 represents *confident* predictions [32]. In the model of monomeric AtCYT6, 70% of the residues had a pLDDT value higher than 70, translating into a confident prediction. Overall, the model displayed two distinct domains with cystatin-like fold, that were connected by an inter-domain linker. Each domain consisted of an α-helix running across an antiparallel 4-stranded β-sheet (Fig. 2A). The N-terminal domain harbored the inhibitory motifs towards PLCPs, and the C-terminal domain contained the legumain binding site. Importantly, when we used AlphaFold to predict a dimeric form of AtCYT6, it predicted the assembly of a DS-dimer AtCYT6 / AtCYT6’ in which the N-terminal region of AtCYT6-NTD of one chain swapped out and integrated at the equivalent position of the other, and *vice versa*. (Fig. 2B). As a consequence of this rearrangement, the PLCP binding sites of both chains were disrupted. The C-terminal domains of AtCYT6 and AtCYT6’ remained with a non-domain-swapped structure and did not interact with each other. In this model 50 % of the residues had a predicted local distance difference test (pLDDT) value higher than 70 [33]. The remaining 50 % of the residues corresponded to the termini, loops, the β5-strands of AtCYT6-CTD and the former L1-loops of the AtCYT6-NTDs. In a next step, we were interested whether computational models of AtCYT6-NTD and AtCYT6-CTD would result in domain-swapped (DS) dimers similarly to AtCYT6. Again, AlphaFold Multimer predicted a DS-dimer of AtCYT6-NTD that comprised the same features of the DS-dimer of AtCYT6 (Fig. S2A). Interestingly, the modeled dimer of AtCYT6-CTD (89 % of its residues with pLDDT value > 80) also assembled via domain swapping with the same topology found for AtCYT6 and AtCYT6-NTD (Fig. S2B). Overall, the models predicted by AlphaFold Multimer reproduced the current model of domain swapping by cystatins.

**Figure 2.**
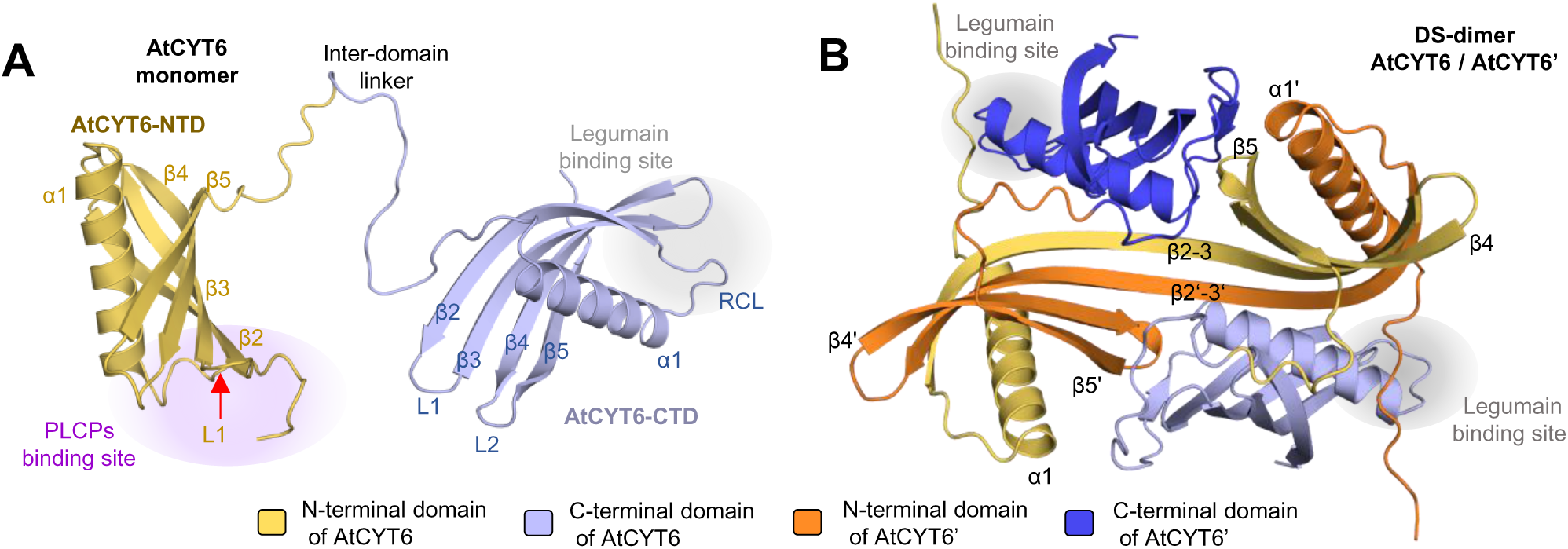
AlphaFold model suggests that dimerization of AtCYT6 may be mediated by domain swapping of AtCYT6-NTD. **(A)** Consistent with our experimental data, the model of AtCYT6 generated by AlphaFold suggested that it is built up by two individual cystatin domains (AtCYT6-NTD and AtCYT6-CTD) that are linked by the inter-domain linker. The binding site for papain-like proteases (PLCPs) is indicated in purple, the L1-loop by a red arrow and the legumain binding site in grey. **(B)** The model predicted by AlphaFold Multimer for the dimer of AtCYT6 (AtCYT6 / AtCYT6’) depicts a domain-swapped dimer (DS-dimer) assembled via the N-terminal domain. In the model the dimer is built up by two AtCYT6 monomers where the N-terminal part of the AtCYT6-NTD and AtCYT6-NTD’ subdomains (β1-α1-β2-L1-) were swapped out and integrated into the equivalent position of AtCYT6-NTD and *vice versa*. Thereby the β2-L1-β3 segment of AtCYT6-NTD (yellow) and AtCYT6-NTD’ (orange) were merged into a single long β2-3 strand, resulting in the loss of the PLCP binding site. In the model, the AtCYT6-CTDs of AtCYT6 (light blue) and AtCYT6’ (dark blue) remained non-domain-swapped and retained the legumain binding site (highlighted in grey). Secondary elements were numbered according to hCC.

### AtCYT6-NTD dimer showed a reduction in AtLEGβ inhibition

In a next step we were wondering whether the formation of AtCYT6 dimers was indeed mediated by domain swapping. To analyze this, we tested AtCYT6 dimers for their ability to inhibit papain. The papain inhibitory site differs in monomeric and dimeric domain-swapped cystatins because the L1 loop, which is essential for PLCP inhibition, is destroyed by the dimer interface. We found that the AtCYT6-NTD dimer was still able to inhibit papain, however at a 10-times higher IC50 compared to monomeric AtCYT6-NTD (Fig. 3). This finding suggested that the AtCYT6 dimers did not - or only partly - form via classical domain-swapping. Thus, AtCYT6-NTD dimers may be composed of (i) non-DS-dimers with reduced affinity towards papain, (ii) a mixture of DS-dimers and non-DS-dimers, or (iii) non-classical swapped dimers where the L1 loop is still intact.

**Figure 3.**
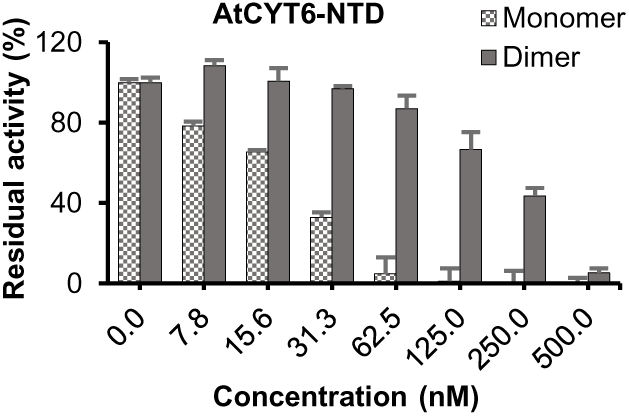
AtCYT6-NTD dimers do not form (exclusively) via domain swapping. Peak fractions of the experiment shown in Figure 1B were analyzed for their ability to inhibit papain in a fluorescence substrate assay. The inhibitory activity of dimeric AtCYT6-NTD towards papain was reduced as compared to the monomeric form. However, AtCYT6-NTD dimers retained their inhibitory activity towards papain, albeit with reduced apparent affinity. This experiment implied that AtCYT6-NTD dimers did not primarily form via domain swapping.

### AtCYT6 forms amyloid fibrils at acidic pH and upon heating

Knowing that AtCYT6 formed high molecular weight oligomers in solution, we were in a next step wondering whether it would also form amyloid fibrils. Given the dramatic structural rearrangements that are typically required to form such fibrils, a considerable amount of ‘activation energy’ must be applied to a fibril-forming protein. To overcome the activation energy barrier, the native conformation can be destabilized by various means, including proteolytic processing, pH shift, and heating to induce fibril formation *in vitro* [12, 23]. In order to access AtCYT6’s overall conformational stability, we monitored its thermal denaturation at different pH values via nano differential scanning fluorimetry (nanoDSF) experiments. Buried aromatic residues (e.g. tryptophan) display a fluorescence peak at 330 nm, and solvent-exposed ones peak at 350 nm. Consequently, the ratio between the fluorescence intensities measured at 350 nm and 330 nm reflects structural rearrangements that result from changes in the chemical environment of aromatic residues. The denaturation curves obtained for AtCYT6 showed destabilization upon pH decrease (Fig. 4A, table 1). The analysis software detected inflection temperatures of 68.5 °C at pH 5.5 and 77.8 °C at pH 7.5 respectively and no inflection point at pH 4.0. Rather than denaturation, we suspect that the observed inflection points corresponded to conformational rearrangements during amyloid fibril formation. To test this hypothesis, we used a thioflavin T (ThT) assay to investigate AtCYT6 fibril formation in a pH and temperature-dependent manner. Indeed, the ThT assay showed that fibrils were formed by AtCYT6 at pH 7.5 upon heating to > 60 °C. At pH 3.0, a readily visible white precipitate was formed already at room temperature, and a strong ThT fluorescence signal was detected at all temperatures tested (Fig. 4B). This observation was in nice agreement with the lack of inflection seen in the thermal denaturation curve of AtCYT6 at pH 4.0, and indicated that fibril formation may have happened already prior to the intrinsic fluorescence analysis. To further confirm the formation of amyloid fibrils, we performed X-ray diffraction experiments. Specifically, we induced fibril formation at 80 °C for 30 minutes at pH 3.0 or 7.5, harvested and dried the samples, and analyzed them via X-ray diffraction experiments. The results depicted in Fig. 4C,D revealed two rings at approximately 4.6 Å and 10 Å resolution, perfectly matching the typical diffraction pattern of cross-beta structures with corresponding β-strand stacking repeats. This experiment confirmed that AtCYT6 indeed formed amyloid fibrils upon conformational destabilization.

**Figure 4.**
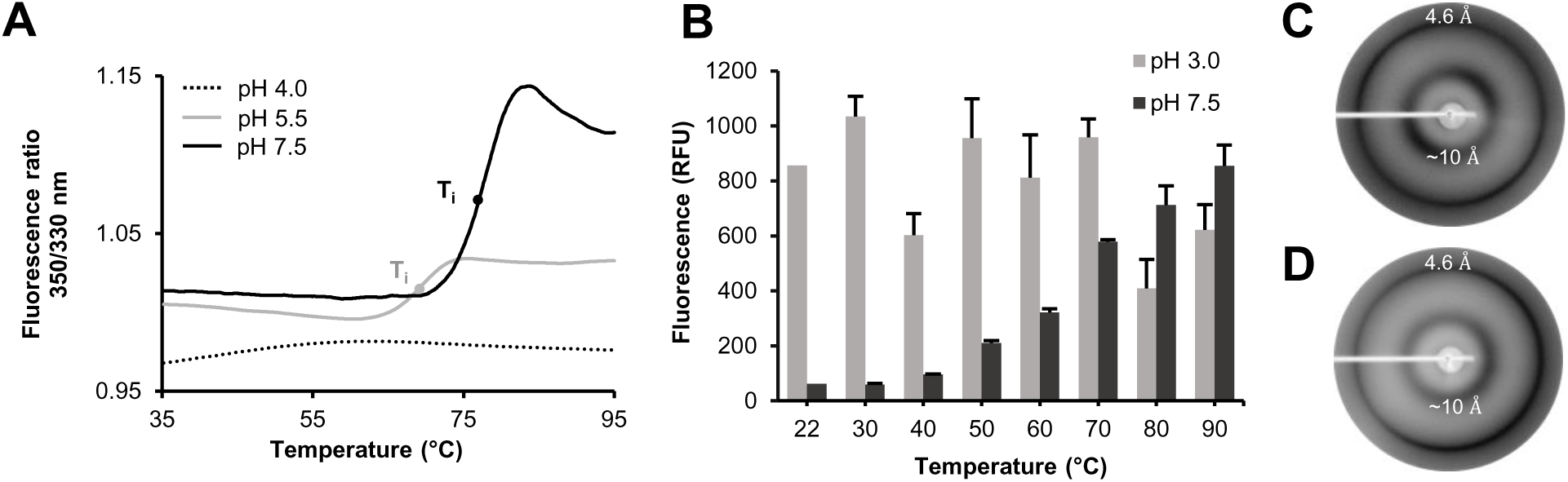
AtCYT6 forms amyloid fibrils upon conformational destabilization. **(A)** Melting curves of AtCYT6 were recorded at indicated pH values via differential scanning fluorimetry (nanoDSF) assays. The denaturation curves obtained at pH 5.5 (grey line) and pH 7.5 (full black line) unveiled unfolding events indicated by inflection temperature (T_i_). No T_i_ was detected at pH 4.0. **(B)** AtCYT6 was incubated at pH 3.0 or 7.5 at indicated temperatures. Subsequently, fibril formation was measured as an increase in thioflavin T fluorescence (ThT) upon binding to the fibrils. While amyloid fibril formation displayed a strong temperature dependency at pH 7.5, a high ThT fluorescence signal was detected in samples incubated at pH 3.0 in all temperatures tested. The X-ray diffraction pattern of fibrils obtained at pH 3.0 **(C)** and pH 7.5 **(D)** after heat treatment displayed two rings at 4.6 Å and approximately 10 Å, revealing the presence of cross-β-sheets characteristic for amyloid fibrils.

**Table 1.** Inflection temperatures detected by nanoDSF measurement at indicated pH values as depicted in Figures 4 and 5.

|  | **AtCYT6** | **AtCYT6-NTD** | **AtCYT6-CTD** |
| --- | --- | --- | --- |
| **pH** | **T<sub>i</sub> (°C)** | **T<sub>i</sub> (°C)** | **T<sub>i</sub> (°C)** |
| **4.0** | - | 42.5 | - |
| **5.5** | 68.5 | 76.2 | 86.2 |
| **7.5** | 77.7 | 80.3 | - |

### AtCYT6-NTD triggers fibril formation

To get mechanistic insights into AtCYT6 fibril formation, we investigated the amyloidogenic potential of the individual N- and C-terminal domains. Along that line, we tested the thermal stability of AtCYT6-NTD and AtCYT6-CTD at different pH values via nanoDSF experiments (Fig. 5A,B). Interestingly, AtCYT6-NTD was drastically destabilized at pH 4.0. An inflection temperature was detected at T_i_ = 42.5 °C. At pH 5.5, AtCYT6-NTD showed a higher thermal stability with an inflection temperature at T_i_ = 76.2 °C and at pH 7.5 an inflection temperature of T_i_ = 80.3 °C was measured (Table 1). Interestingly, we observed a second conformational transition at pH 4.0 and 5.5 at approx. 10 °C above the first transition. In line with these observations, a ThT test of AtCYT6-NTD showed that it formed amyloid fibrils at pH 3.0 already at temperatures > 40 °C and at pH 7.5 at temperatures > 80 °C (Fig. 5C). The X-ray diffraction pattern of the amyloid fibrils obtained from heat-treated AtCYT6-NTD further confirmed its amyloid nature (Fig. 5E). This thermal stability profile was in stark contrast to AtCYT6-CTD, which did not show an increase in ThT fluorescence upon pH shift or heating and no significant inflection temperatures in nanoDSF experiments (Fig. 5D, table 1). Importantly, despite the data pointing towards a remarkable overall stability of AtCYT6-CTD, we did observe AtCYT6-CTD fibril formation at pH 3.0 in a single occasion, which was however not reproducible. Fibril formation was confirmed by a ThT assay and X-ray diffraction experiments (Supplementary Fig. S3A,B). That indicated that although AtCYT6-NTD possesses the predominant amyloidogenic potential, AtCYT6-CTD is in principle also able to form amyloid fibrils, although only under very strict conditions.

**Figure 5.**
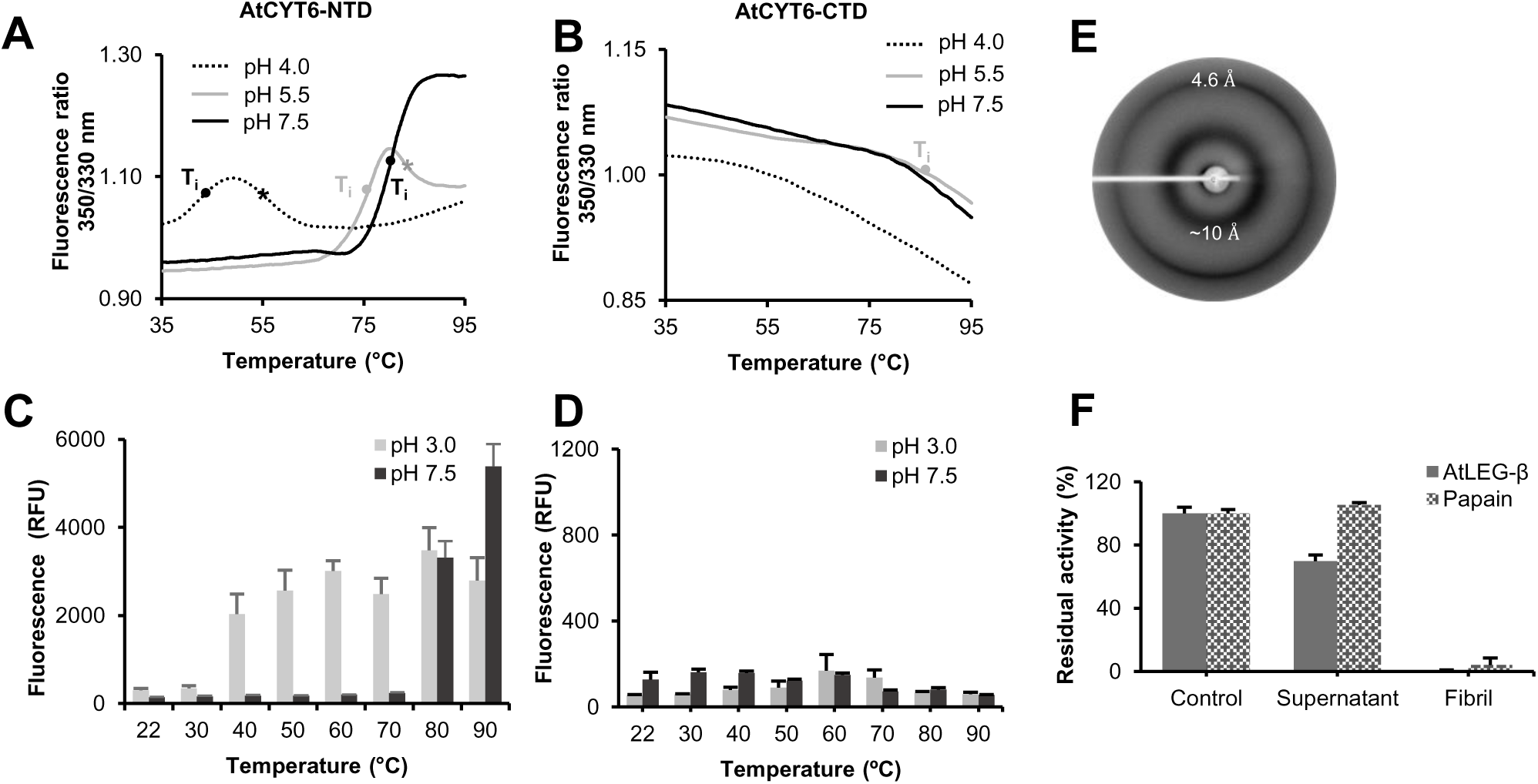
AtCYT6-NTD promotes amyloid fibril formation. Differential scanning fluorimetry (nanoDSF) experiments of AtCYT6-NTD **(A)** and AtCYT6-CTD **(B)** showed that similar to AtCYT6, AtCYT6-NTD displayed pH-dependent inflection temperatures. A second conformation transition was observed at pH 4.0 and 5.5 (labeled with a star). No T_i_ was detected for AtCYT6-CTD, except at pH 5.5 above 80 °C, suggesting an overall high thermal stability. AtCYT6-NTD **(C)** and AtCYT6-CTD **(D)** were incubated at pH 3.0 or 7.5 at indicated temperatures. Subsequently, fibril formation was measured as an increase of thioflavin T (ThT) fluorescence upon binding to the fibrils. Fibrils were formed by AtCYT6-NTD at pH 3.0 from 40 °C on, whereas at pH 7.5 an increase in fluorescence was only detectable at 80 °C and above. AtCYT6-CTD did not yield amyloid fibrils at both pHs and all temperatures. **(E)** X-ray diffraction experiments of AtCYT6-NTD fibrils obtained upon incubation at pH 3.0 and 80 °C revealed two diffraction rings at 4.6 Å and approximately 10 Å resolution, characteristic for cross-β-sheet structures. **(F)** AtCYT6 fibrils were washed to remove residual amounts of soluble protein until their supernatant did not significantly inhibit the enzymes compared to the control (buffer). The residual activity of papain (white dotted columns) and legumain (grey columns) in the presence of buffer, supernatant or the fibrils was measured. The fibrils strongly inhibited both enzymes, revealing the presence of functional inhibitors.

### AtCYT6 fibrils inhibit papain and legumain

In a next step we were wondering whether AtCYT6 amyloid fibrils contained functional inhibitors. If domain swapping was a prerequisite for fibril formation, we expected that AtCYT6 fibrils would lose their ability to inhibit PLCPs. To test this, we analyzed the inhibition of AtLEGβ and papain by AtCYT6 fibrils after several cycles of washing to eliminate soluble inhibitors (Fig. 5F). Surprisingly, the insoluble fibrils conserved the ability to inhibit both AtLEGβ and papain, implying that AtCYT6 fibrils contained functional inhibitor proteins. This finding is inconsistent with the classical domain swapping mechanism where the PLCP-inhibitory L1-loop is incorporated into an extended β2-3 strand. Similarly, the observed non-covalent AtCYT6-NTD dimer could also not be sufficiently explained by classical domain swapping. Therefore, instead we suggest that fibrils form via a slightly modified swapping-mechanism, e.g. where only the α-helix but not the β2 strand swap and repositions itself into another AtCYT6 molecule. In this helix-only swapping mode (Fig. 6), the L1-loop stays intact and could therefore explain the functional PLCP-inhibition by the AtCYT6-NTD dimers and the fibrils. Additionally, this swapping mode could also explain why the dimer is unstable and short lived, since it harbors fewer stabilizing interactions as compared to the classical dimer swapping mode. An alternative explanation may be that the fibrils form via classical domain-swapping and be decorated with co-precipitated native AtCYT6 protein. However, this mechanism does not explain why dimeric AtCYT6-NTD was still a functional PLCP inhibitor.

**Figure 6.**
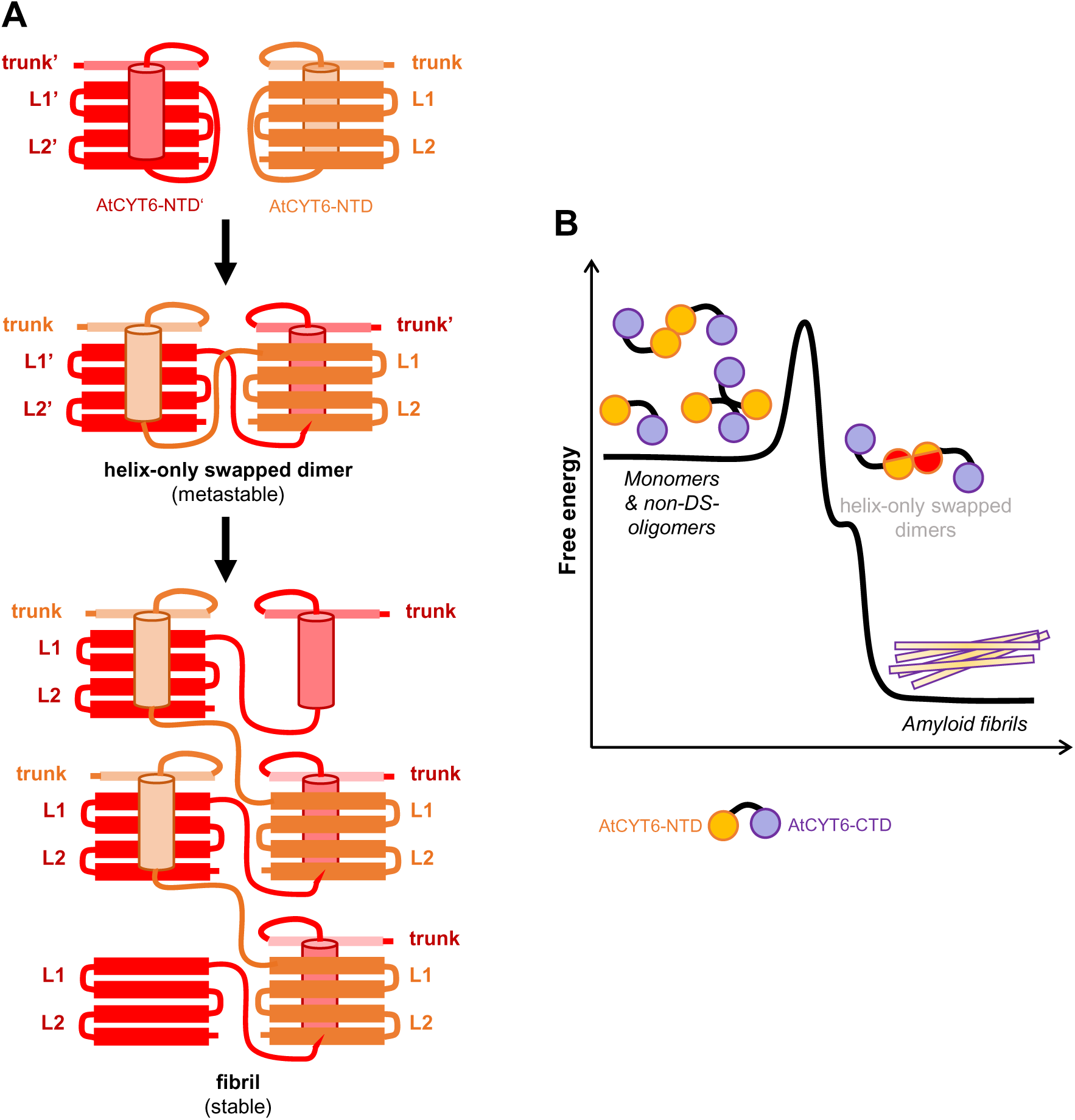
Conversion of AtCYT6 to amyloid fibrils requires energy. **(A)** Model of the helix-only swapping mode of AtCYT6-NTD. In this model, a metastable dimer state is rapidly converted to stable fibrils. **(B) F**ree energy model suggesting that amyloid fibrils are the energetically preferred oligomerization state of AtCYT6. To reach the fibril state, monomeric and non-domain swapped dimers and oligomers (non-DS-oligomers) need to overcome a certain energy barrier to convert to short-lived, metastable helix-only swapped dimers and subsequently to fibrils.

## Discussion

The cystatin superfamily comprises multiple single domain animal cystatins known to form amyloid fibrils *in vitro* and *in vivo* [9, 10, 12]. Our work conveys compelling evidence of fibril formation *in vitro* by a multidomain cystatin. These findings allow us to postulate that amyloid fibril formation is a conserved feature of cystatin-like proteins across kingdoms and different sub-families. The current model of fibril formation by cystatins proposes dimerization via domain swapping prior to the assembly of fibrils [8]. In this study we show that AtCYT6 is present in different oligomeric states in solution, with both covalent and non-covalent dimers. Covalent dimer formation was mediated by disulfide formation of Cys138 whereas non-covalent dimer formation could be attributed to interactions of AtCYT6-NTD. Since the non-covalent dimer was still able to inhibit papain, it can be excluded that the dimer formed exclusively via classical domain swapping. The L1-loop, which is a critical part of the PLCP binding site would be disrupted in the classic DS-dimer [12, 34, 35]. However, instead we suggest that AtCYT6-NTD dimers are composed of non-DS-dimers with reduced affinity towards papain, or a mixture of DS-dimers and non-DS-dimers, or non-classical swapped dimers like e.g. helix-only swapped dimers where the L1 loop is still intact. Within this study we were not able to isolate a stable, DS-dimer of AtCYT6, which might be explained by the thermal stability profile of AtCYT6. In a previous study we found that the monomer - DS-dimer transition of hCE and hCC takes place at 65 °C and fibril formation was observed at > 90 °C at pH 5.5 (Δ 25 °C) [12]. Similarly, AtCYT6 showed two transitions in nanoDSF experiments at pH 5 (Fig. 5A). However, while the two transitions were separated by approx. 25 °C in hCE and hCC, they were apart by < 10 °C in AtCYT6. Since the two transitions were close, they likely represent overlapping equilibria. Based on this observation, we propose that similar as in hCE and hCC, the first transition temperature corresponds to conformational rearrangements that precede amyloidogenesis, e.g. helix swapping. Since the proposed helix-only swapped dimer is less stable and therefore only short lived, it may readily convert into more stable fibrils (Fig. 6B). Furthermore, fibril formation was triggered already at a lower temperature in AtCYT6 as compared to hCE, which is also consistent with the suggested helix-only swapped dimer since it is less stable than a classical DS-dimer.

Within this study we found that AtCYT6-NTD is mainly responsible for the amyloidogenesis of AtCYT6. Oligomerization and amyloidogenesis of hCC has been broadly studied in the context of hereditary cystatin C amyloid angiopathy (HCCAA), where it gives rise to amyloid deposits in human tissues [11, 36, 37]. The L1 hinge loop of hCC harbors the Q-I-V-A-G sequence containing the Q-X-V-X-G motif, a key element for PLCPs inhibition. Importantly, this motif is integrated into the β2-3 strand upon domain swapping [24]. Rodziewicz-Motowidło and colleagues analyzed 11 cystatin structures deposited to the PDB and identified distorted dihedral *ψ* and *φ* angles of the glycine and valine residues on their hinge loops [38], and single point-mutations within this region enabled the modulation of the monomer/DS-dimer ratio [39]. Furthermore, non-domain-swapping proteins engineered to bear the hinge loop sequence of stefin B (Q-V-V-A-G) in solvent-exposed β-turns were successfully driven to domain swapping, illustrating the relevance of this sequence in the dimerization of cystatins [40]. Sequence alignments between DS-dimer-forming cystatins hCE, hCC, and phycys from *H. lupulus*, *V. unguiculata*, *S. officinarum* and the individual domains of AtCYT6 (Supplementary Fig. S4) revealed that while AtCYT6-NTD harbored the conserved sequence motif, AtCYT6-CTD did not. This observation is in nice agreement with our experimental data that showed that AtCYT6-CTD was not a PLCP inhibitor, and that AtCYT6-NTD enables dimerization and fibril formation [22].

Finally, we suggest that AtCYT6 fibrils might possess a core composed of domain swapped AtCYT6-NTDs that expose to the solvent monomeric C-terminal domains covalently attached to the fibril via the interdomain linker. As fibrils retained inhibitory capacity against papain and AtLEGβ, we propose that they may form via a modified swapping mode e.g. by helix-only swapping or co-precipitate with functional inhibitors, as previously observed in hCE fibrils [12]. Fibrils could provide a stationary stabilization “docking port” for AtLEGβ extracellularly, as AtCYT6-CTD has been shown to stabilize AtLEGβ from pH-driven denaturation under non-acidic conditions [22]. We furthermore found evidence that AtCYT6 was present as separate AtCYT6-NTD and AtCYT6-CTD subdomains in planta, most likely induced by proteolytic processing [22]. Importantly, this observation suggests that fibrils may be formed by AtCYT6-NTD only, without association of AtCYT6-CTD (and *vice versa*). Other regulators of amyloidogenesis have been identified and are translatable to the plant cell environment, such as temperature, proteolytic processing/truncation, glycosylation, point mutations, presence of divalent cations and combinations of these factors [10, 12, 23, 39, 41–44].

### Material & Methods

#### AtCYT6 constructs design

The AtCYT6 constructs AtCYT6 full-length, -NTD and -CTD were designed as described previously [22]. Briefly, the sequence encoding AtCYT6 and lacking the N-terminal signal peptide (M1-M34) was based on the sequence of cystatin 6 from *A. thaliana* found on Uniprot under the accession code Q8HOX6-2. It was cloned into the pET-15b vector and therefore AtCYT6 harbored an N-terminal His_6_-tag, as well as AtCYT6-CTD and AtCYT6-CTD_long_. The AtCYT6-NTD sequence was cloned into the pET-28a vector and the final construct contained a C-terminal His_6_-tag.

#### AtCYT6 expression and purification

Proteins were expressed and purified following the procedure previously described [22]. AtCYT6, -NTD, -CTD and -CTD_long_, as well as their single-point mutants, were expressed in *E. coli* BL21 (DE3) cells in the presence of ampicillin (for constructs in pET15b vector) or kanamycin (pET-28a) in 500 mL of LB medium in 2 L expression flasks shaking at 230 rpm. The cultures were pre-inoculated and grown at 37 °C until OD_600_ reached 0.8-1.0. The expression was induced with IPTG at a final concentration of 1 mM and carried out overnight at 25 °C. On the next day, cells were harvested by centrifugation, lysed by sonication and subsequently centrifuged for 1 h at 4 °C and 17500 g to sediment insoluble cell components. The soluble fractions of the proteins of interest were purified by immobilized metal affinity chromatography (IMAC) using Ni^2+^-beads. The resin was equilibrated in a buffer containing 50 mM Tris pH 7.5 and 300 mM NaCl. After the lysates were loaded on the beads, they were washed with increasing amounts of imidazole (0 mM, 5 mM, 10 mM and 20 mM) in equilibration buffer. Proteins were eluted with 250 mM imidazole in the same buffer. Further purification was carried out using gel filtration chromatography. Specifically, an Äkta FPLC system (Cytiva) equipped with a superdex 75 (S75) or superdex 200 (S200) column equilibrated in a buffer composed of 20 mM Tris pH 7.5 and 50 mM NaCl (SEC buffer) was used.

#### Preparation of AtLEGβ and papain

Papain from *Carica papaya* was purchased from Merck (Darmstadt, Germany). Preparation of AtLEGβ included the heterologous expression of the enzymatically inactive proAtLEGβ in *Leishmania tarentolae* cells (LEXSY, Jena Bioscience) purification of the proenzyme from the supernatant of culture media using IMAC (Ni^2+^-beads), and pH-driven self-activation following the protocols previously described [45].

#### Oligomerization state evaluation by size exclusion chromatography

For size exclusion chromatography experiments with AtCYT6, -NTD, -CTD and -CTD_long_, proteins were concentrated up to 2 mg/ml in a buffer composed of 20 mM Tris pH 7.5 and 50 mM NaCl (SEC buffer). For constructs harboring a cysteine residue, i.e. AtCYT6 and AtCYT6-CTD_long_, DTT (dithiothreitol) was added at 2 mM final concentration to reduce disulfide bonds when desired. 500 µl of protein sample were injected into a S200 (in the case of AtCYT6) or S75 (AtCYT6-CTD, -NTD and CTD_long_) column equilibrated in SEC buffer and coupled to an Äkta FPLC system (Cytiva). Experiments were done at room temperature.

To test the effect of pH on the oligomerization state of AtCYT6, we repeated the same experiment using buffers composed of 20 mM Tris pH 7.0, 50 mM NaCl and 2 mM DTT or 20 mM citric acid pH 5.5, 50 mM NaCl and 2 mM DTT. The influence of ionic strength on the oligomerization state was evaluated using buffers composed of 20 mM Tris pH 7.5, 2 mM DTT and either 50 mM or 500 mM NaCl. To test the stability of different oligomeric species, we reinjected the respective peak fractions using SEC buffer supplemented with 2 mM DTT. The protein samples in all the assay conditions described above consisted of 500 μL of AtCYT6 solution at 2 mg/mL injected into a S200 column connected to an Äkta FPLC system (Cytiva) at room temperature. The effect on the oligomerization state of a 5-fold dilution of AtCYT6 was evaluated in SEC buffer with 2 mM DTT upon injection of 500 μL of AtCYT6 solution at either 0.4 mg/mL or 2.0 mg/mL in the same setup.

#### Cross-linking with glutaraldehyde

Proteins were buffer exchanged into 20 mM citric acid pH 5.5 and 100 mM NaCl to a final concentration of 0.3 mg/mL and incubated with freshly prepared glutaraldehyde solution at a final concentration of 0.175 % (w/v) or buffer (control) at room temperature. Samples were taken after 1, 5 and 10 minutes and transferred to new tubes containing 1 μL of 2.5 M Tris-HCl pH 8.8 to stop the reaction. Subsequently, samples were analyzed by SDS-PAGE.

#### Modeling of AtCYT6 monomer and dimers

The sequence we used to model monomeric AtCYT6 was lacking the N-terminal signal peptide and comprised residues ranging from Ala35 to Asp234, according to Uniprot ID Q8H0X6. Modeling of the monomer was performed using AlphaFold. Dimeric AtCYT6 was modeled using Alphafold Multimer via the Colab notebook using two copies of the AtCYT6 sequence. Likewise, dimer models of AtCYT6-NTD (Ala35 – Asp127) and AtCYT6-CTD (Gly141 – Asp234) were obtained using two copies of the respective sequence as input. Models were illustrated with PyMOL Molecular Graphics System (Schrödinger, LLC).

#### Differential scanning fluorimetry (nanoDSF)

To access the thermal stability of different AtCYT6 constructs, we performed differential scanning fluorimetry experiments. The proteins to be analyzed were incubated in assay buffer composed of 20 mM Tris 7.5 and 50 mM NaCl, or 20 mM citric acid pH 5.5 or 4.0 and 50 mM NaCl for 5 minutes at a final protein concentration of 1 mg/ml. For constructs harboring a cysteine residue, DTT (dithiothreitol) was added to the assay buffer at a final concentration of 2 mM. Suitable capillaries were filled with protein solution (approximately 10 μL) and changes in intrinsic fluorescence intensity were measured at 330 and 350 nm using a Tycho NT.6 instrument (Nanotemper) upon heating the samples from 35 °C to 95 °C within a total measuring time of approx. 3 minutes.

#### Thioflavin T assay

Proteins in SEC buffer were buffer exchanged and concentrated to 10 mg/mL in a buffer composed of 50 mM citric acid pH 3.0, 100 mM NaCl and 2 mM DTT if suitable, or 50 mM Tris pH 7.0, 100 mM NaCl and 2 mM DTT if suitable. 10 µl of the target protein were transferred to PCR tubes and loaded into a thermocycler (Eppendorf, Hamburg, Germany). Subsequently, proteins were incubated for 30 min at 30, 40, 50, 60, 70, 80 or 90 °C, immediately followed by cooling to 4 °C for 20 min. The samples incubated at 22 °C were not transferred to the thermocycler, but instead were kept at room temperature for 30 min and then put on ice for 20 min. For the fluorescence assay with thioflavin T (ThT), a stock solution of ThT (Sigma-Aldrich) was prepared by dissolving 8 mg of ThT in 10 ml of PBS buffer. The ThT stock was diluted in a 1:50 ratio in PBS, and 49 μL were pipetted into a 384-well black polystyrene non-binding surface plate (Corning, Kennebunk, USA) followed by the addition of 1 μL of thoroughly resuspended protein sample. Four replicates for each condition were measured. The signal was monitored in a Tecan M200 plate reader (Tecan) with absorption at 440 nm and emission at 482 nm.

#### X-ray diffraction experiments

For X-ray diffraction experiments, proteins (10 mg/ml) were heated to 80 °C for 30 min and immediately afterwards incubated on ice for 20 min. Precipitates were collected by centrifugation, the excess supernatant was removed, and the pellet was dried using a Speedvac (Eppendorf, Hamburg, Germany). The remainder was a plastic-like dried piece of precipitate, which could be mounted onto the tip of a quartz glass capillary. X-ray diffraction was assayed in house using a Bruker Microstar rotating anode generator mounted with a Mar345dtb detector.

#### Inhibition of AtLEGβ and papain by AtCYT6 fibrils

The fibril suspension used in this inhibition assay was obtained by heating AtCYT6 at a concentration of 10 mg/mL at 80 °C for 30 minutes in a buffer composed of 50 mM citric acid pH 3.0, 100 mM NaCl and 2 mM DTT, followed by cooling for 20 minutes on ice. The fibrils were subsequently washed to eliminate soluble proteins until the UV_280_ of the supernatant was null, and more than 80 % of enzymatic activity was recovered (compared to buffer as control). The washing consisted of multiple cycles of centrifugation at 13000 g for 1 min, elimination of the supernatant, and resuspension in 1 mL of fresh buffer. For the measurement of the residual activity of AtLEGβ in the presence of fibrils, 1 μL of washed fibril suspension was added to 44 μL of the synthetic peptidic substrate Z-Ala-Ala-Asn-AMC (Bachem) diluted in reaction buffer (20 mM citric acid pH 5.5, 100 mM NaCl, 0.02 % Tween 20, and 2 mM DTT) at a final concentration of 40 μM. The reaction was started with the addition of 5 μL of active enzyme at 500 nM concentration diluted in reaction buffer. The increase in fluorescence (in RFU) over time was measured by an Infinite M200 Plate Reader (Tecan) at 25 °C using UV-STAR 96-well microplates (Greiner, Kremsmünster, Austria). As controls, the measurement of AtLEGβ activity was performed in the presence of 1 μL of the supernatant of the fibril suspension, and 1 μL of buffer 50 mM citric acid pH 3.0, 100 mM NaCl and 2 mM DTT. Four replicates of each condition were measured. The exact same protocol was used for the assessment of papain activity in the presence of AtCYT6 fibrils, except that we used the synthetic peptidic substrate Z-Phe-Arg-AMC (Bachem), and the reaction buffer was composed of 20 mM MES pH 6.5, 50 mM NaCl, 0.02 % Tween 20, and 2 mM DTT.

#### Sequence alignments

A multiple sequence alignment was prepared using CLUSTAL O (1.2.4) using as input the full-length sequences of human cystatin C (hCC; Uniprot P01034), human cystatin E (hCE; Uniprot Q15828), canecystatin-1 (Uniprot Q7Y0Q9), cystatin 1 from hop (sequence retrieved from PDB 6VLQ), cystatin 1 from cowpea (Uniprot A0A1X9Q255) and cystatin 6 from *A. thaliana* (Uniprot Q8H0X6).

## Acknowledgements

The authors wish to thank Martina Wiesbauer for technical assistance.

## Funding and additional information

This work was supported by the Austrian Science Fund (FWF, project number P31867, to E.D.).

