## Supporting Information for "*Arabidopsis thaliana* phytocystatin 6 forms functional oligomer and amyloid fibril states"

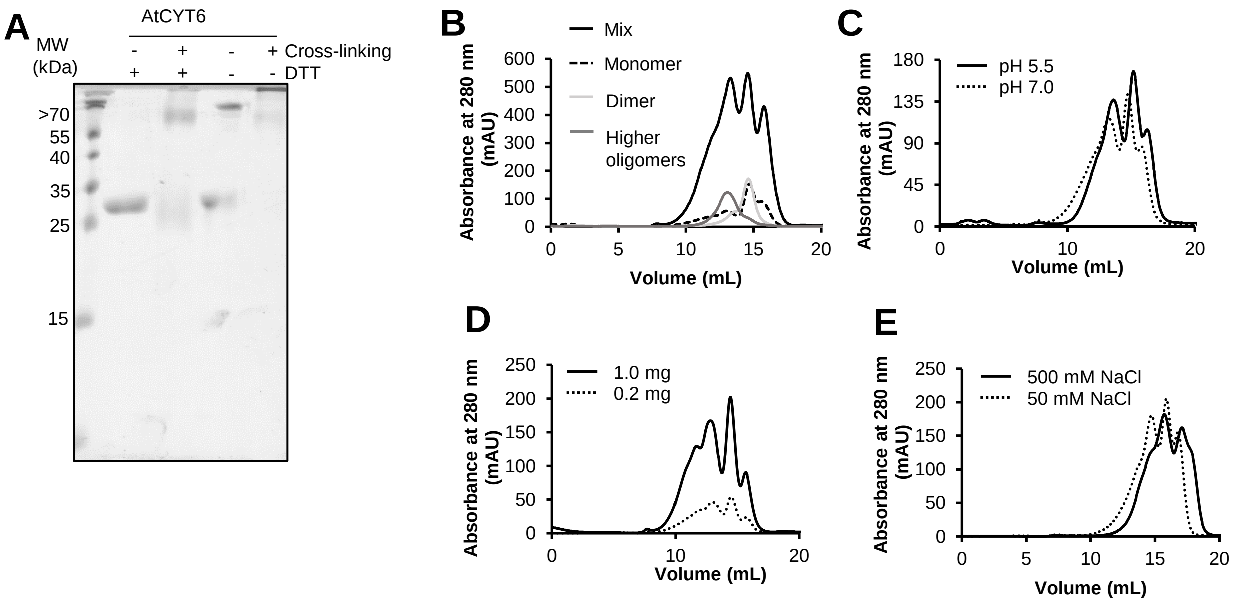


**Supplementary Figure 1. AtCYT6 exists in different oligomeric states.**

**(A)** Glutaraldehyde cross-linking of reduced and non-reduced AtCYT6 showed that the presence of DTT (dithiothreitol) increased the subpopulation of monomers and dimers in solution. **(B)** Size exclusion chromatography (SEC) experiments with AtCYT6 (black line) under reducing conditions showed that upon reinjection of separately collected peaks of monomers (dashed black line), dimers (light grey line) and higher oligomers (dark grey line) into the chromatography column, the monomers were unstable and tended to convert into dimers and higher oligomers, whereas dimers and higher oligomers were oligomerization states relatively stable in solution. **(C)** SEC experiments of AtCYT6 at pH 5.5 or pH 7.5 under reducing conditions revealed pH-independent oligomerization. Oligomerization was similarly observed at different protein concentrations **(D)** and ionic strengths **(E)**.

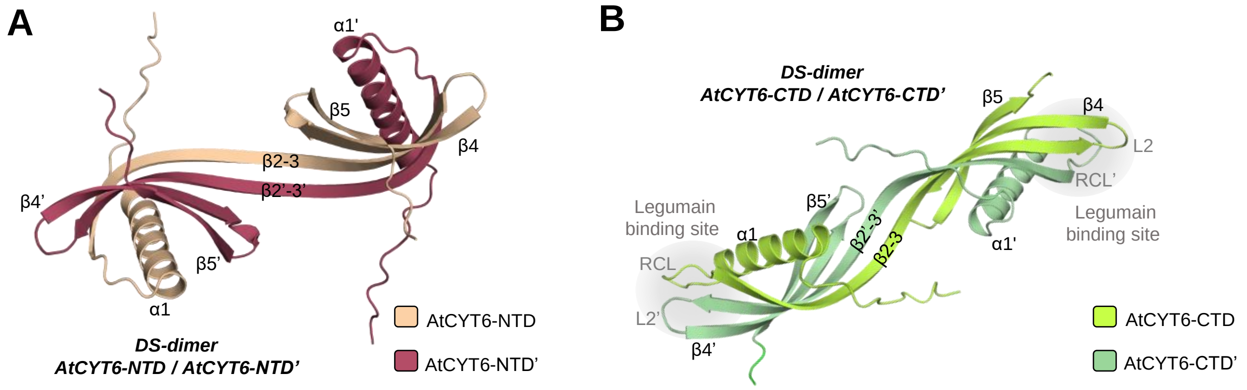


**Supplementary Figure 2. Molecular modeling suggests that AtCYT6-NTD and AtCYT6-CTD may form dimers via domain-swapping.**

AlphaFold Multimer predicted dimerization via domain-swapping in **(A)** AtCYT6-NTD and **(B)** AtCYT6-CTD. The models suggest that the N-terminal β1-α1-β2-L1-β3 segments of AtCYT6-NTD (wheat) and AtCYT6-NTD' (red) swap out and reposition themselves into the complementary molecule. The N-terminal segments of AtCYT6-CTD (light green) and AtCYT6-CTD' (dark green) swap in a similar way.

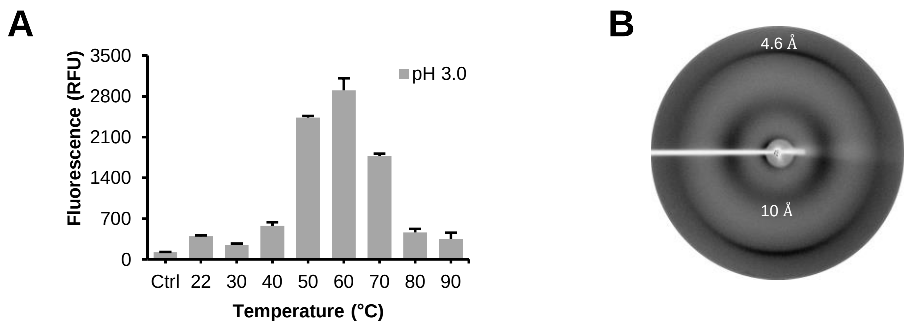


**Supplementary Figure 3. AtCYT6-CTD formed amyloid fibrils in a single occasion.**

**(A)** The thioflavin T (ThT) fluorescence assay indicated amyloid fibril formation at pH 3.0 upon heat treatment from 50 to 70 ºC. **(B)** X-ray diffraction experiment of the fibrils obtained in (A) revealed the two diffraction maxima at 4.6 Å and 10 Å resolution characteristic for cross-beta structures.

```
CLUSTAL O(1.2.4) multiple sequence alignment

hCC            MA---GPLRAPLLLLAILAVALAVSPAAGSSPGKPPRLVGGPMDASVE---EEGVRRALD
hCE            MARSNLPLALGL---ALVAFCLLALPR-DARARPQERMVGELRDLSPD---DPQVQKAAQ
AtCYT6CTD      -----------------------------------GEHESGWREVPGD---DPEVKHVAE
SoCYT1         --------------------------MAEAHNGRRVGMVGDVRDAPAGHENDLEAIELAR
AtCYT6NTD      MMRSRFLLFIVFFSLSLF--ISSLIASDLGFCNEEMALVGGVGDV-PANQNSGEVESLAR
HlCYT1         -----------------------------------MATVGGIKEV-DGNQNSLEIESLAR
VuCYT1         -----------------------------------MAALGGNRDV-AGNQNSLEIDSLAR
                                                      .   :       .

hCC            FAVGEYNKASNDMYHSRALQVVRARKQIVAGV-NYFLDVELGRTTCTKTQPN-----LDN
hCE            AAVASYNMGSNSIYYFRDTHIIKAQSQLVAGI-KYFLTMEMGSTDCRKTRVTGDHVDLTT
AtCYT6CTD      QAVKTIQQRSNSLFPYELLEVVHAKAEVTGEAAKYNMLLKLKR-GEKE------------
SoCYT1         FAVAEHNSKTNAMLEFERL--VKVRHQVVAGT-MHHFTVQVKEAGGGK------------
AtCYT6NTD      FAVDEHNKKENALLEFARV--VKAKEQVVAGT-LHHLTLEILE-AGQK------------
HlCYT1         YAVDEHNKKQNSLLQFEKV--VNTKQQVVSGT-IYIITLEAVD-GGKK------------
VuCYT1         FAVEEHNKKQNALLEFGRV--VSAQQQVVSGT-LYTITLEAKD-GGQK------------
                **   :   * :        : .: ::..    : : ::       :

hCC            CPFHDQPHLKRKAFCSFQIYAVPWQGTMTLSKSTCQDA------- 146
hCE            CPLAAGA-QQEKLRCDFEVLVVPWQNSSQLLKHNCVQM------- 149
AtCYT6CTD      ------------EKFKVEVHKNH-EGALHLNHAEQHHD------- 94
SoCYT1         ------------KLYEAKVWEKVWENFKQLQSFQPVGDA------ 106
AtCYT6NTD      ------------KLYEAKVWVKPWLNFKELQEFKPASD------- 127
HlCYT1         ------------KVYEAKVWEKPWMNFKELQEFKLIGDAPSGSSA 101
VuCYT1         ------------KVYEAKVWEKPWLNFKELQEFKHVGDAPA---- 97
                              . ::      .   *
```

**Supplementary Figure 4. Sequence alignments indicate that the Q-X-V-X-G motif is a critical determinant of domain swapping.**

The sequences of AtCYT6-NTD and AtCYT6-CTD were aligned with sequences of cystatins where crystal structures of domain swapped dimers were available. The alignment was created using sequences of human cystatin C (hCC; Uniprot P01034), human cystatin M/E (hCE; Uniprot Q15828), canecystatin-1 (SoCYT1; Uniprot Q7Y0Q9), cystatin 1 from hop (HlCYT1; sequence retrieved from PDB 6VLQ), cystatin 1 from cowpea (VuCYT1; Uniprot A0A1X9Q255) and cystatin 6 from *A. thaliana* (Uniprot Q8H0X6, split to AtCYT6-NTD and AtCYT6-CTD). The Q-X-V-X-G motif critical for PLCP inhibition and present in the hinge loop L1 (black box) is conserved amongst all analyzed sequences, except AtCYT6-CTD.
